# Inside Out: the physiology of *Brucella* Vegetative and Intracellular Growth

**DOI:** 10.1101/2024.08.06.606627

**Authors:** Nazarena Wade, Diego J. Comerci, Alfonso Soler-Bistué, María Inés Marchesini

## Abstract

Growth rate is a key prokaryotic trait that allows estimating fitness and understanding cell metabolism. While in some clades it has been well studied in model organisms, there is little data on slow-growing bacteria. In particular, there is a lack of quantitative studies on the species of the *Brucella* genus. This is an important microorganism since it is the causative agent of brucellosis, one of the most widespread bacterial zoonoses affecting several species of animals, including humans. *Brucella* species exhibit approximately 97% genomic similarity. Despite this, *Brucella* species show different host preferences, zoonotic risks, and pathogenicity. After more than one hundred years of research, numerous aspects of *Brucella* biology, such as *in vivo* and *in vitro* growth remain poorly characterized. In this work, we analyzed vegetative and intracellular growth of the classical *Brucella* species (*B. abortus* 2308, *B. melitensis* 16M. *B. suis* 1330, *B. ovis*, and *B. canis*). Strikingly, each species displayed particular growth parameters in culture. Doubling time (DT) spanned from 2.7 hs^-1^ in *B. suis* to 18h^-1^ for *B. ovis*. In the context of intracellular infection of J774A.1 phagocytic cells, DT was higher, but it widely varied across species, closely correlating to the growth observed *in vitro*. Overall, despite high similarity of the genomes, we found species-specific growth parameters in the intracellular cell cycle.

## Introduction

Duplication is one of the main functions of life. The cell cycle is the key process that, at the cellular level, reproducibly allows the cells to properly divide their DNA, generating enough cell components to give rise to two daughter cells. At the population level, most lifeforms follow a logistic growth. In single celled organisms, particularly in prokaryotes, population growth is measured using the growth curve, a method that has been a Microbiology workhorse for more than a century[1]. The doubling time (DT) of any culturable species can be obtained directly from the exponential phase of the growth[2]. More recently, methods arose for non-culturable bacteria[3]. DT is a central trait for prokaryotes since it allows to quantify its duplication capacity while it permits estimating its fitness and understanding cell metabolism and biochemistry[1, 4]. DT has taxonomic value presenting a high phylogenetic inertia, in other words, it is highly conserved among closely related species[5, 6].

Recently, bioinformatic approaches relying only on genome sequence allowed predicting DTs with great precision[6, 7]. We now know that DT can widely vary across prokaryotes ranging from a few minutes to months or even years[8]. Overall, there is a link between genome structure, the DT of a given organism and its ecological niche and function[9-11]. For instance, factors such as codon usage, ribosomal RNA operon number, the quantity of tRNAs encoded in a genome and the genomic location of transcription and translation genes may help predict DT[6, 7, 12, 13]. However, the experimental determination of DT remains limited and a great majority of what we know of cell physiology is limited to well-studied models such as *Escherichia coli, Bacillus subtilis* or *Caulobacter crescentus*. Unculturable or slow growing microorganisms remain much less studied.

Meanwhile, while the DT reflects the maximum growth capacity and is mostly measured *in vitro*, microorganisms face different circumstances according to their ecological niche[9, 10]. Moreover, many microorganisms have complex life cycles that include different stages. The DT may vary in each one of them or, in other words, in some the parts of their life cycle, growth may be below the minimum DT.

Bacteria from a-Proteobacterota are microorganisms of high interest since many of them establish close interactions with multicellular hosts. It includes biotechnologically relevant microorganisms such as *Rhizobiaceae*, well-known as crop symbionts. Also, important pathogens such as *Agrobacterium* and *Brucella* [14]. This last genus encompasses a growing array of bacteria traditionally linked to brucellosis in major livestock species, as well as zoonotic infections in humans[15].

*Brucella* species exhibit a wide host range, historically recognized through six “classical” species named after their preferred hosts: *Brucella abortus* (cattle), *Brucella melitensis* (goats and sheep), *Brucella ovis* (sheep), *Brucella suis* (pigs), *Brucella canis* (dogs), and *Brucella neotomae* (desert rats)[16]. The genus has expanded during in the last years however, *Brucella* species share a remarkably low genetic diversity, with approximately 97% similarity between species [17, 18].

Brucellosis, a neglected and re-emerging zoonotic disease, is globally distributed. Among human infections, *B. melitensis* poses the greatest pathogenic risk, followed by *B. abortus* and *B. suis*, while *B. canis* presents a lower zoonotic potential[19]. Transmission primarily occurs through contact with infected animal secretions, ingestion, or aerosols, with raw milk consumption and occupational exposure as significant risk factors [20, 21]. *Brucella* growth kinetics *in vitro* and *in vivo* is key to understanding brucellosis.

The detailed study of *Brucella* physiology is challenging due to their relatively slow growth, their selective agent status accompanied with their high infectivity trough aerosols. At the population level, there are very few studies precisely measuring their physiological parameters[22, 23]. Growth curve studies, following optical density over time, allow determination of key physiological parameters such as growth rate (or its inverse, the DT), the lag phase duration and the maximum system carrying capacity. Performing growth curves in *Brucella* is particularly difficult since it implies long-term exposure to frequent culture handling. As a consequence, *Brucella* growth curves and physiological parameters remain poorly characterized throughout the bibliography. However, a better understanding of its physiology will necessarily contribute to our understanding of the mechanisms underlying host preference and virulence. In parallel, infection of cell systems by *Brucella* is well characterized. After cell entry, *Brucella* load is drastically reduced due to interactions with the degradative pathway. However, a subset of cells persists and starts intracellular replication. Therefore, during infection experiments, reduction in intracellular bacteria loads is observed until 8-10 hs post-infection. Then, intracellular replication occurs increasing bacterial loads to at least two orders of magnitude. Finally, *Brucella* exploits multivesicular bodies to exit host cells[24]. Although *Brucella* intracellular stages have been extensively characterized, studies quantifying intracellular DT are lacking. Curiously, several studies have addressed some aspects of *Brucella* cell cycle both in culture and in the context of cell infection[25-28] but population studies are lacking.

In this work, we deepen into the study of population growth kinetics of five classical and representative *Brucella* species: *B. abortus* 2308, *B. melitensis* 16M. *B. suis* 1330, *B. ovis*, and *B. canis*. We characterized their *in vitro* physiology at the population level using automatized growth curves, a method that allows running many growth curves in parallel in diverse growth conditions that allow safe manipulation. In addition, since *Brucella* species are facultative intracellular pathogens with limited environmental survival that rely on interactions with various host cell types to establish infection, we decided to compare and correlate vegetative to intracellular growth rates. For this, we performed intracellular replication assays at different time points that allowed estimating growth rates in the context of infection. Interestingly, we found 4-fold differences in growth parameters despite their phylogenetic proximity.

## Materials and Methods

### Bacterial strains and culture conditions

*Brucella* strains were cultured on Tryptic Soy Agar, *Brucella* Broth (BB), or Tryptic Soy Broth (TSB) at 37°C. Liquid cultures were grown for 16-24 hs under agitation (200 rpm), except for *B. ovis*. Culture mediums were supplemented with nalidixic acid (5 µg/ml). All manipulations involving *Brucella* species were performed at the biosafety level 3 (BSL3) laboratory facility at the Universidad Nacional de San Martín.

### Genome comparisons across *Brucella* species

For genome comparisons we employed two independent programs to perform pair-wise calculation of the average nucleotide identity (ANI) the program ANI calculator by Kostas lab and OrthoANIu tool described elsewhere[29, 30]. Both approaches provided equivalent results.

### Automatic growth curve measurements

For automated growth curve measurements, 96-well plates were utilized, avoiding the use of external rows and columns. Overnight cultures were diluted 1/1000 in BB and distributed in triplicates in 96-well microplates. Growth curve experiments were conducted using a TECAN plate reader, measuring OD_620nm_every 15 minutes at 37°C with maximum agitation, except for *Brucella ovis*, where no agitation was used. Growth parameters including growth rate and lag phase were determined using QurvE software analysis[31].

### Intracellular replication assays

For the standard antibiotic protection assay, murine macrophage-like J774A.1 cells were seeded in 24-well plates at a density of 10^5^ cells per ml and incubated overnight at 37°C before the infection. *Brucella* strains, grown in TSB or BB with nalidixic acid for 24 hs, were diluted in culture medium prior to infection. The bacterial suspension was added to achieve a multiplicity of infection of 100:1 and centrifuged at 1500 rpm for 10 minutes. After 30 minutes of incubation at 37°C, cells were washed, and fresh medium containing streptomycin (100 µg/ml) and gentamicin (50 µg/ml) was added. At 4, 24, 30, 42 and, 48 hours post-infection, cells were washed, lysed, and intracellular colony-forming units (CFU) were determined by direct plating on TSB agar plates.

### Intracellular growth rates calculations

For constructing a growth curve and determining growth rates, the number of generations was calculated using the growth formula with initial and final CFU counts: N(t) = N_0_X 2t/DT. From this, the doubling time (DT) was derived.

Additionally, these calculations were verified using an online growth rate calculator.

## Results

### Determination of *Brucella* growth parameters

Since there is little data on precise growth parameters on *Brucella* species, we performed automated growth curves on the classical species: *B. suis, B. melitenis* 16M, *B. abortus* 2308, *B. ovis* and *B. canis*. For this, strains were grown in Brucella Broth (BB) and OD _620nm_was monitored over time. A representative growth curve is shown in Figure 1a. *Brucella* species showed distinct growth parameters which are summarized in Table 1. *B. suis* was the species displaying the lowest doubling time (2.772 ± 0.275 hs) and the species achieving the highest OD_600_value (Fig. 1a and 1b). *B. melitensis* and *B. canis* behaved similarly, with *B. canis* reaching a higher OD_600_value (Fig. 1a). The doubling time was similar in *B. canis* (3.312 ± 0.096 hs) and *B. melitensis* (3.388 ± 0.159 hs), while the lag phase was longer in *B. canis* (Fig. 1b and 1c). *B. abortus* showed an average doubling time of 4.741 ± 0.164 hs and a lag phase comparable to the lag phase of *B. canis* and, *B. ovis* (Fig. 1b and 1c). Among the species analyzed, *B. ovis* was the slowest growing with a doubling time of 18.140 ± 0.767 hs. However, the lag phase duration was similar to *B. canis* and *B. abortus* (Fig. 1b and 1c). Overall, we observed a variation of up 6-fold in DT between the species when comparing *B. suis* to *B. ovis*. Meanwhile, the duration of lag phases also showed discrepancy but to a lower scale, varying only by two-fold between species. In general, species displaying higher DTs displayed longer durations of lag phases (Fig 1b and 1c). Interestingly, although the species employed in this study display a high genome similarity, reflected by an average nucleotide identity higher than 99% (Table 2), physiological differences are large. The variation of DTs contrasts with the high genomic similarity and the high phylogenetic inertia of this trait.

**Table 1.** Physiological parameters of *Brucella* species in BB medium.

| Strain | Lag Phase (hours) |  |  |  | Generation Time (hours) |  |  |  |
| --- | --- | --- | --- | --- | --- | --- | --- | --- |
|  | Average | Median | SD | N | Average | Median | SD | N |
| *B. suis* 1330 | 13,16 | 13,62 | 1.822 | 10 | 2.772 | 2.774 | 0.275 | 10 |
| *B. abortus* 2308 | 21,18 | 20.22 | 2.156 | 10 | 4.741 | 4.748 | 0.164 | 10 |
| *B. melitensis* 16M | 15,46 | 15.96 | 1.896 | 10 | 3.388 | 3.346 | 0.159 | 10 |
| *B. canis* | 14.22 | 15.01 | 3.470 | 5 | 3.312 | 3.295 | 0.096 | 5 |
| *B. ovis* | 25,23 | 26.51 | 2.293 | 5 | 18.14 | 17.95 | 0.767 | 5 |

**Table 2.** Average Nucleotide Identity (ANI) between representative *Brucella* species.

| ANI | *B. ovis* | *B. canis* | *B. melitensis*<br>16M | *B. abortus*<br>2308 | *B. suis* 1330 |
| --- | --- | --- | --- | --- | --- |
| *B. suis* 1330 | 99.67 | 99.9 | 99.67 | 99.69 | 100 |
| *B. abortus*<br>2308 | 99.59 | 99.69 | 99.9 | 100 |  |
| *B. melitensis*<br>16M | 99.60 | 99.69 | 100 |  |  |
| *B. canis* | 99.62 | 100 |  |  |  |
| *B. ovis* | 100 |  |  |  |  |

**Figure 1.**
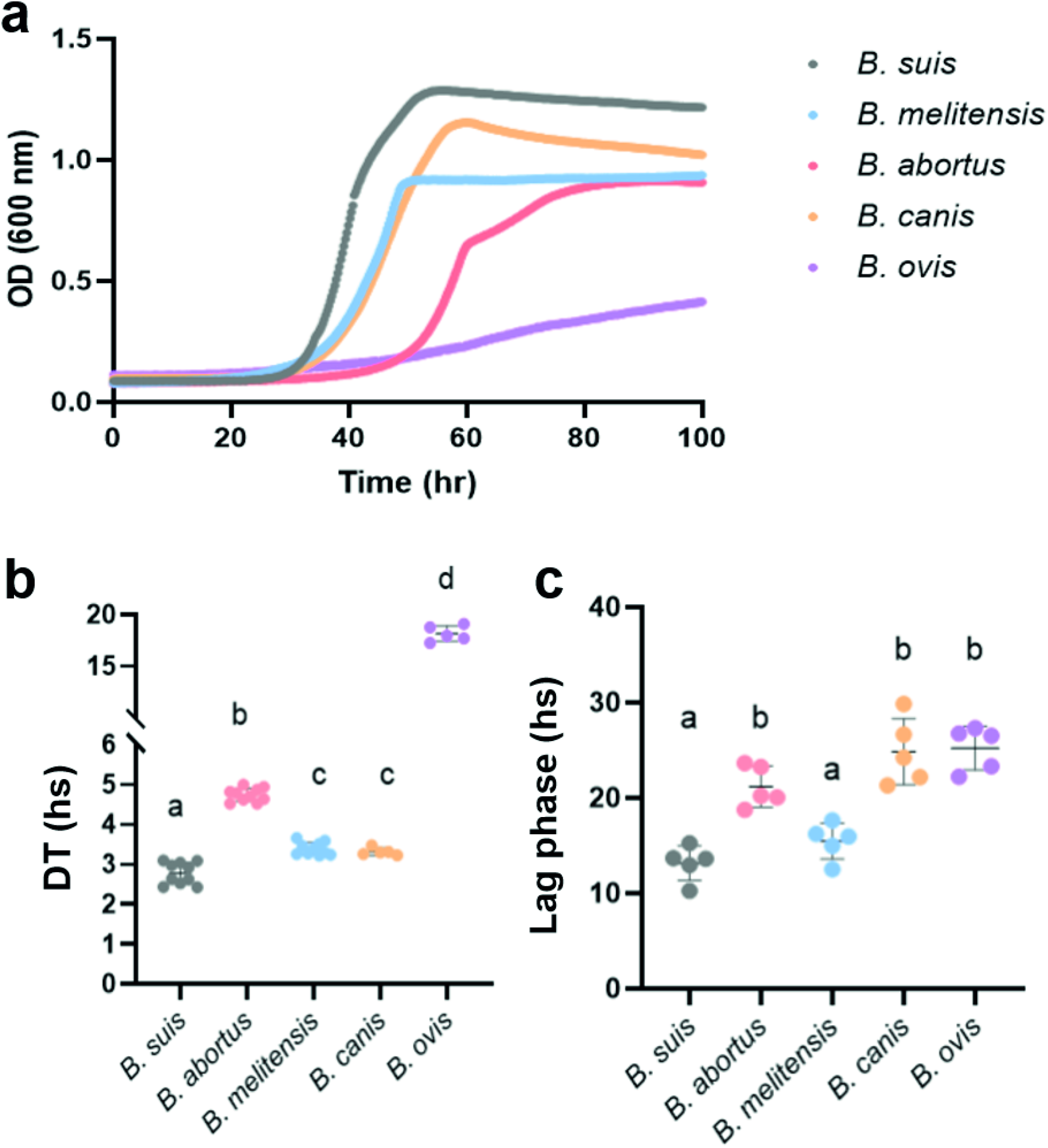
Growth parameters differ between Brucella species. **(a)** Representative growth curves of the depicted *Brucella* species in BB medium. **(b)** Doubling times (DT) and **(c)** Lag phase duration was calculated from automated growth curves in BB medium. Individual values and the means are shown. Statistical significance was analyzed using One-way ANOVA followed by Tukey for multiple comparisons. Different letters denote significant differences between strains (*P < 0.0001).

### *Brucella* intracellular growth quantification

*Brucella* intracellular life cycle is complex and highly coordinated with the host. Bacteria adhere and invade host cells, subvert lysosomal degradation, replicate within specialized organelles, and ultimately exit to restart the infectious cycle. In this section, the invasion and intracellular replication profiles of the selected species were analyzed and compared. For this, bacterial loads were determined using a gentamicin protection assay in macrophagic-like cells of the cell line J774A.1. Cells were infected as described in methods, and at different times post-infection (p.i.) the cells were lysed and the colony forming units (CFU) enumerated (Fig. 2a). The five species showed no statistically significant differences at 4 hs p.i., and retrospective CFU counts showed that cells were infected with an equivalent number of bacteria of each species. These results demonstrate that the invasion capacity of the species studied is similar in J774A.1 cells. At 24 hs p.i., when intracellular replication had already begun, the species with the lowest CFU/ml values were *B. abortus* and *B. ovis*. At 30 hs p.i., this trend was maintained and cells infected with *B. suis, B. melitensis* and *B. canis* showed the highest bacterial loads. Later, when the replicative niche is well established within cells, *B. suis* and *B. canis* were the species with the highest intracellular CFU/ml values, while *B. abortus* and *B. melitensis* showed intermediate values. Throughout the experiment, the cells infected with *B. ovis* contained less CFU/ml than the cells infected with the other *Brucella* species. Together, these results indicate that intracellular replication rates in macrophagic-like cells differ among species.

**Figure 2.**
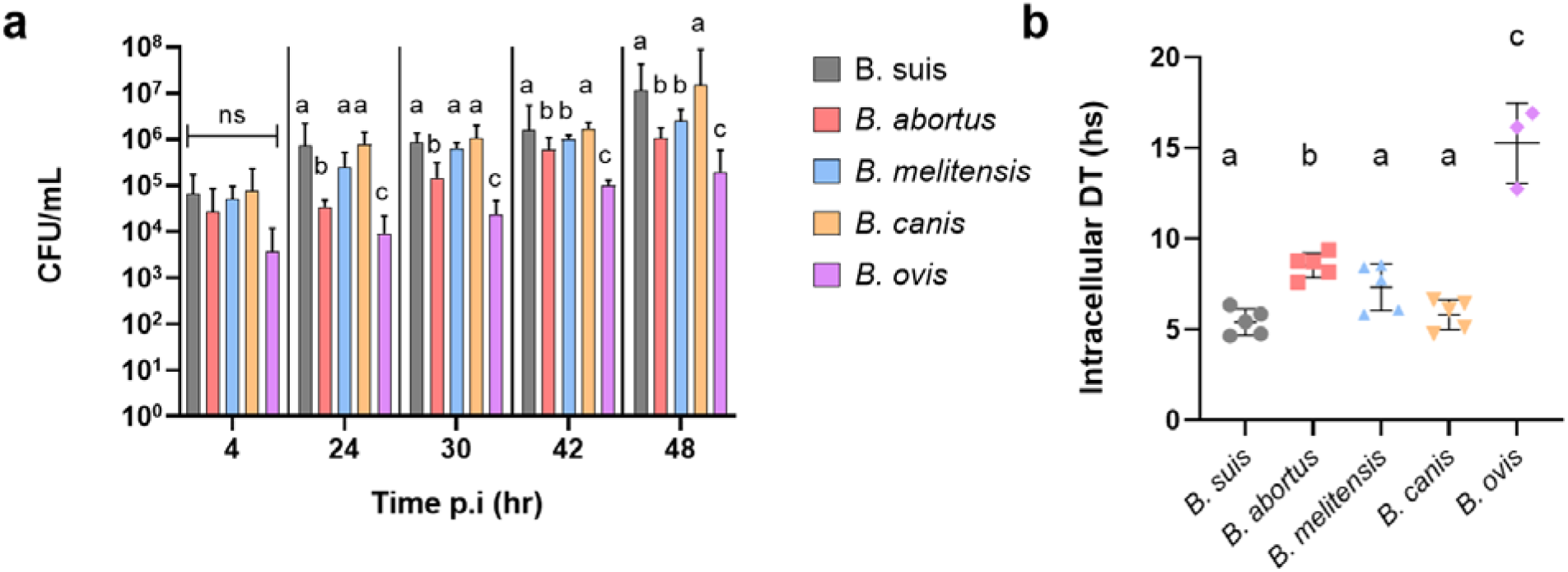
Brucella intracellular growth. **(a)** Intracellular replication in J774A.1 cells of the indicated *Brucella* species. Statistical significance was analyzed using Two-way ANOVA followed by Tukey for multiple comparisons. Different letters indicate significant differences between strains (One-way ANOVA) **(b)** Intracellular DTs were calculated from gentamicin protection assays as explained in the methods section. Statistical significance was analyzed using One-way ANOVA followed by Tukey for multiple comparisons. Different letters indicate significant differences between strains (One-way ANOVA *P < 0.0001).

To compare the extracellular and intracellular multiplication rates of the studied *Brucella* species, we calculated the intracellular DTs as described in the methods section. Results displayed Figure 2b and Table 3, show similar trends as in the previous point, with *B. ovis* being the slowest species and *B. suis* the fastest. However, intracellular generational times are longer than vegetative DTs, except for *B. ovis* that replicates at a slower rate outside host cells.

**Table 3.** Growth of *Brucella* species in the intracellular environment.

| Strain | Intracellular Doubling Time<br>(hours) |  |  |  |
| --- | --- | --- | --- | --- |
|  | Average | Median | SD | N |
| *B. suis* 1330 | 5.42 | 5,450 | 0,7200 | 5 |
| *B. abortus* 2308 | 8.54 | 8,740 | 0,6743 | 5 |
| *B. melitensis*<br>16M | 7.34 | 7,739 | 1,288 | 5 |
| *B. canis* | 5.82 | 6,092 | 0,8185 | 5 |
| *B. ovis* | 15.28 | 16,15 | 2,215 | 3 |

## Discussion

Over the past decades, *Brucella* studies have provided substantial insights into infection biology, immunity and host cell biology. However, the basis of *Brucella* host restriction and differential virulence remains poorly understood[32]. *B. abortus* and *B. melitensis* differ in approximately 2400 SNPs, while *B. canis* and *B. suis* biovar 4 have only 253 SNPs differences[33]. Despite this, these species have distinctive host preferences and pathogenicity[34]. Growth capacity of *Brucella* genus has been assessed mostly qualitatively. In this work, we quantified differences in growth parameters, particularly in lag phase duration and DT both during vegetative growth (Figure 1) and in the intracellular niche (Figure 2). Overall, we found that growth during vegetative growth positively correlates to DTs in the context of intracellular replication (r=0.97, p=0.0045). Despite a resemblance of more than 99% according to ANI (Table 2), *B. suis* grew 6.7 times faster than *B. ovis* and twice faster than *B. abortus*. Also, it was 20% faster than *B. melitensis* and *B. canis*. There is a recent reports of a fast-growing *Brucella* species, B. *microtii*. But as in previous works, assessment of growth has been made qualitatively and there is no information of the detailed physiological parameters. The genome analysis of this species indicated that some crucial mutations occur at the level of ribosomal RNA genes (*rrn*)[35]. Indeed, the number and the genomic location of *rrn* correlates with growth rate in most bacteria[6], including related α -Proteobacteria[36, 37]. On the other hand, *B. ovis* also offers a genomic insight explaining the slow growth observed. Degradation of *B*.*ovis* genome, due to a high number of inactivating mutations and transposable elements, would explain growth rate deficency[38]. Indeed, it was demonstrated that the incapability to assimilate CO_2_under the low CO_2_tension of a standard atmosphere is determined by single-nucleotide insertions in the carbonic anhydrase gene, *bcaA[39]*.

## Conclusion

To date, the information on physiological parameters of *Brucella* species was very limited and growth was inferred from colony size or the time needed for colonies to appear in agar plates. By using high throughput methods that ensure safe manipulation, we were able to contribute to determining with high precision the DTs and the lag phase duration of five representative species of the genus. We also assessed the growth rate capacity in the intracellular milieu. We found that, while genomes are highly similar, the physiological parameters may differ widely among the strains both *in vitro* and during macrophage infection. This is somehow surprising in a trait such as growth with high phylogenetic inertia. Future works should assess these few genomic differences that impact in the DTs of *Brucella* species, which in turn could be useful to understand genomic factors that shape bacterial growth and virulence.

## References

1. Jun, S., et al., Fundamental principles in bacterial physiology-history, recent progress, and the future with focus on cell size control: a review. Rep Prog Phys, 2018. 81(5): p. 056601.

2. Monod, J., The Growth of Bacterial Cultures. Annual Reviews in Microbiology, 1949. 3: p. 371–394.

3. Emiola, A. and J. Oh, High throughput in situ metagenomic measurement of bacterial replication at ultra-low sequencing coverage. Nat Commun, 2018. 9(1): p. 4956.

4. Schaechter, M., A brief history of bacterial growth physiology. Front Microbiol, 2015. 6: p. 289.

5. Walkup, J., et al., The predictive power of phylogeny on growth rates in soil bacterial communities. ISME Commun, 2023. 3(1): p. 73.

6. Vieira-Silva, S. and E.P. Rocha, The systemic imprint of growth and its uses in ecological (meta)genomics. PLoS Genet, 2010. 6(1): p. e1000808.

7. Weissman, J.L., S. Hou, and J.A. Fuhrman, Estimating maximal microbial growth rates from cultures, metagenomes, and single cells via codon usage patterns. Proc Natl Acad Sci U S A, 2021. 118(12).

8. Madin, J.S., et al., A synthesis of bacterial and archaeal phenotypic trait data. Sci Data, 2020. 7(1): p. 170.

9. Roller, B.R., S.F. Stoddard, and T.M. Schmidt, Exploiting rRNA operon copy number to investigate bacterial reproductive strategies. Nat Microbiol, 2016. 1(11): p. 16160.

10. Couso, L.L., et al., Ecology theory disentangles microbial dichotomies. Environ Microbiol, 2023. 25(12): p. 3052–3063.

11. Soler-Bistue, A., L.L. Couso, and I.E. Sanchez, The evolving copiotrophic/oligotrophic dichotomy: From Winogradsky to physiology and genomics. Environ Microbiol, 2023.

12. Rocha, E.P., Codon usage bias from tRNA’s point of view: redundancy, specialization, and efficient decoding for translation optimization. Genome Res, 2004. 14(11): p. 2279–86.

13. Hu, X.P. and M.J. Lercher, An optimal growth law for RNA composition and its partial implementation through ribosomal and tRNA gene locations in bacterial genomes. PLoS Genet, 2021. 17(11): p. e1009939.

14. Batut, J., S.G. Andersson, and D. O’Callaghan, The evolution of chronic infection strategies in the alpha-proteobacteria. Nat Rev Microbiol, 2004. 2(12): p. 933–45.

15. Godfroid, J., et al., Brucellosis at the animal/ecosystem/human interface at the beginning of the 21st century. Prev Vet Med, 2011. 102(2): p. 118–31.

16. Moreno, E., A. Cloeckaert, and I. Moriyon, Brucella evolution and taxonomy. Vet Microbiol, 2002. 90(1-4): p. 209–27.

17. Tsolis, R.M., Comparative genome analysis of the alpha-proteobacteria: relationships between plant and animal pathogens and host specificity. Proc Natl Acad Sci U S A, 2002. 99(20): p. 12503–5.

18. Moreno, E., The one hundred year journey of the genus Brucella (Meyer and Shaw 1920). FEMS Microbiol Rev, 2021. 45(1).

19. Godfroid, J., et al., From the discovery of the Malta fever’s agent to the discovery of a marine mammal reservoir, brucellosis has continuously been a re-emerging zoonosis. Vet Res, 2005. 36(3): p. 313–26.

20. Blasco, J.M. and B. Molina-Flores, Control and eradication of Brucella melitensis infection in sheep and goats. Vet Clin North Am Food Anim Pract, 2011. 27(1): p. 95–104.

21. Xavier, M.N., et al., Pathological, immunohistochemical and bacteriological study of tissues and milk of cows and fetuses experimentally infected with Brucella abortus. J Comp Pathol, 2009. 140(2-3): p. 149–57.

22. Del Giudice, M.G., et al., PhiA, a Peptidoglycan Hydrolase Inhibitor of Brucella Involved in the Virulence Process. Infect Immun, 2019. 87(8).

23. Hauschild, A.H. and H. Pivnick, Continuous culture of Brucella abortus S.19. Can J Microbiol, 1961. 7: p. 491–505.

24. Marchesini, M.I., J.M. Spera, and D.J. Comerci, The ‘ins and outs’ of Brucella intracellular journey. Curr Opin Microbiol, 2024. 78: p. 102427.

25. Brown, P.J., et al., Polar growth in the Alphaproteobacterial order Rhizobiales. Proc Natl Acad Sci U S A, 2012. 109(5): p. 1697–701.

26. Deghelt, M., J.J. Letesson, and X. De Bolle, On the link between cell cycle and infection of the Alphaproteobacterium Brucella abortus. Microb Cell, 2014. 1(10): p. 346–348.

27. Deghelt, M., et al., G1-arrested newborn cells are the predominant infectious form of the pathogen Brucella abortus. Nat Commun, 2014. 5: p. 4366.

28. Vassen, V., et al., Localized incorporation of outer membrane components in the pathogen Brucella abortus. EMBO J, 2019. 38(5).

29. Goris, J., et al., DNA-DNA hybridization values and their relationship to whole-genome sequence similarities. Int J Syst Evol Microbiol, 2007. 57(Pt 1): p. 81–91.

30. Yoon, S.H., et al., A large-scale evaluation of algorithms to calculate average nucleotide identity. Antonie Van Leeuwenhoek, 2017. 110(10): p. 1281–1286.

31. Wirth, N.T., et al., QurvE: user-friendly software for the analysis of biological growth and fluorescence data. Nat Protoc, 2023. 18(8): p. 2401–2403.

32. O’Callaghan, D. and A.M. Whatmore, Brucella genomics as we enter the multi-genome era. Brief Funct Genomics, 2011. 10(6): p. 334–41.

33. Foster, J.T., et al., Whole-genome-based phylogeny and divergence of the genus Brucella. J Bacteriol, 2009. 191(8): p. 2864–70.

34. Suarez-Esquivel, M., et al., Brucella Genomics: Macro and Micro Evolution. Int J Mol Sci, 2020. 21(20).

35. Audic, S., et al., Brucella microti: the genome sequence of an emerging pathogen. BMC Genomics, 2009. 10: p. 352.

36. Medici, I.F., et al., The distinct cell physiology of Bradyrhizobium at the population and cellular level. BMC Microbiol, 2024. in press.

37. Cherni, A.E. and X. Perret, Deletion of rRNA Operons of Sinorhizobium fredii Strain NGR234 and Impact on Symbiosis With Legumes. Front Microbiol, 2019. 10: p. 154.

38. Tsolis, R.M., et al., Genome degradation in Brucella ovis corresponds with narrowing of its host range and tissue tropism. PLoS One, 2009. 4(5): p. e5519.

39. Varesio, L.M., et al., A Carbonic Anhydrase Pseudogene Sensitizes Select Brucella Lineages to Low CO(2) Tension. J Bacteriol, 2019. 201(22).

